# Instability of cooperation in finite populations

**DOI:** 10.1101/707927

**Authors:** Chai Molina, David J. D. Earn

## Abstract

Evolutionary game theory has been developed primarily under the implicit assumption of an infinite population. We rigorously analyze a standard model for the evolution of cooperation (the multi-player snowdrift game) and show that in many situations in which there is a cooperative evolutionarily stable strategy (ESS) if the population is *infinite*, there is no cooperative ESS if the population is *finite* (no matter how large). In these cases, contributing *nothing* is a globally convergently stable finite-population ESS, implying that apparent evolution of cooperation in such games is an artifact of the infinite population approximation. The key issue is that if the size of groups that play the game exceeds a critical proportion of the population then the infinite-population approximation predicts the wrong evolutionary outcome (in addition, the critical proportion itself depends on the population size). Our results are robust to the underlying selection process.

## 1 Introduction

Many evolutionary games assume—for mathematical convenience—that populations are infinitely large (*e.g.*, (1–7)). This assumption is sometimes justified on the grounds that “[p]opulations which stay numerically small quickly go extinct by chance fluctuations” (8, §2.1). Of course, all real populations are finite, and important differences in evolutionary dynamics between finite and infinite populations have been demonstrated (9–15). In spite of the technical challenges of working with finite populations, some exact analytical results have been obtained for two-player games with discrete strategy sets (9, 12, 14–16). However, most existing finite-population results rely on approximation methods and simulations (11, 15, 17–21). Notably, almost all finite-population results involve discrete strategy sets, such as when individuals must choose between making a fixed positive contribution to a public good, or nothing at all (*e.g.*, (9, 12, 14–16)). Yet, evolutionary games involving continuous strategy sets (*e.g.*, allocating time or effort to a communal task) are both widely applicable and extensively studied using infinite-population models (22). Moreover, to our knowledge, all existing results for finite populations depend on a choice of selection process (*e.g.*, Moran or Wright-Fisher (23, 24)).

Here, we present mathematically rigorous results that identify critical differences in the predictions of evolutionary games in finite and infinite populations. We focus on a standard model for exploring the evolution of cooperation—the continuous multi-player snowdrift game (3)—which has previously been studied in infinite populations using exact analysis and simulations (3, 7, 25–27) and in finite populations using approximations and simulations (11, 21, 28, 29).

We show that evolutionary outcomes in finite and infinite populations can be dramatically different. In particular, for a class of snowdrift games for which a cooperative ESS exists in infinite populations (30), we find conditions under which there is no cooperative ESS when played in finite populations. This qualitative difference in predictions for finite and infinite populations can occur no matter how large the finite population is, and is universal in the sense that it is independent of the selection process (31). To our knowledge, there are no other examples in the literature of qualitatively different dynamics in finite and infinite populations that persist for arbitrarily large populations and are independent of the selection process; other studies that demonstrate such differences (*e.g.*, (32)) are restricted to particular selection processes.

The results we present are supported by formal mathematical theorems, which we state in *Results* and prove in *Methods* and *Supporting Information*.

## 2 Terminology

The ***snowdrift game*** is an abstraction of the situation in which a group of individuals encounters a snowdrift that blocks their path. We suppose that *n* players are drawn from a population of self-interested individuals (*n* is the ***group size***), and that each player chooses how much to contribute to a public good—*e.g.*, snow cleared off the path—from which all group members benefit. A focal individual contributing *x* incurs a cost *C*(*x*) that depends only on its own contribution, whereas its benefit *B*(*τ*) depends on the ***total good*** *τ* contributed by the group as a whole. The focal individual’s ***payoff***—which is interpreted as a change in fitness—is then

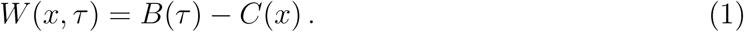

If *x* is a continuous variable, as we assume here, the game is said to be continuous. Positive contributions represent ***cooperative strategies***, and individuals who contribute nothing are said to ***defect***. If the population is finite and contains *N* individuals, then for convenience we refer to the ratio *G* = *N/n* as the ***number of groups***; however, we *do not* assume that the population is simultaneously subdivided into groups of *n* individuals (and in particular, *G* need not be an integer).

To avoid mathematical complexities that are not relevant to the biological issues that concern us, we impose a few natural conditions on the cost and benefit functions and refer to the ***natural snowdrift game*** (NSG; see *Methods §5.1*). The NSG was introduced in (30), where it was shown that—when played in infinite populations—the game always has a cooperative ESS. Cost, benefit and fitness functions for an NSG example are shown in figure 1.

**Figure 1:**
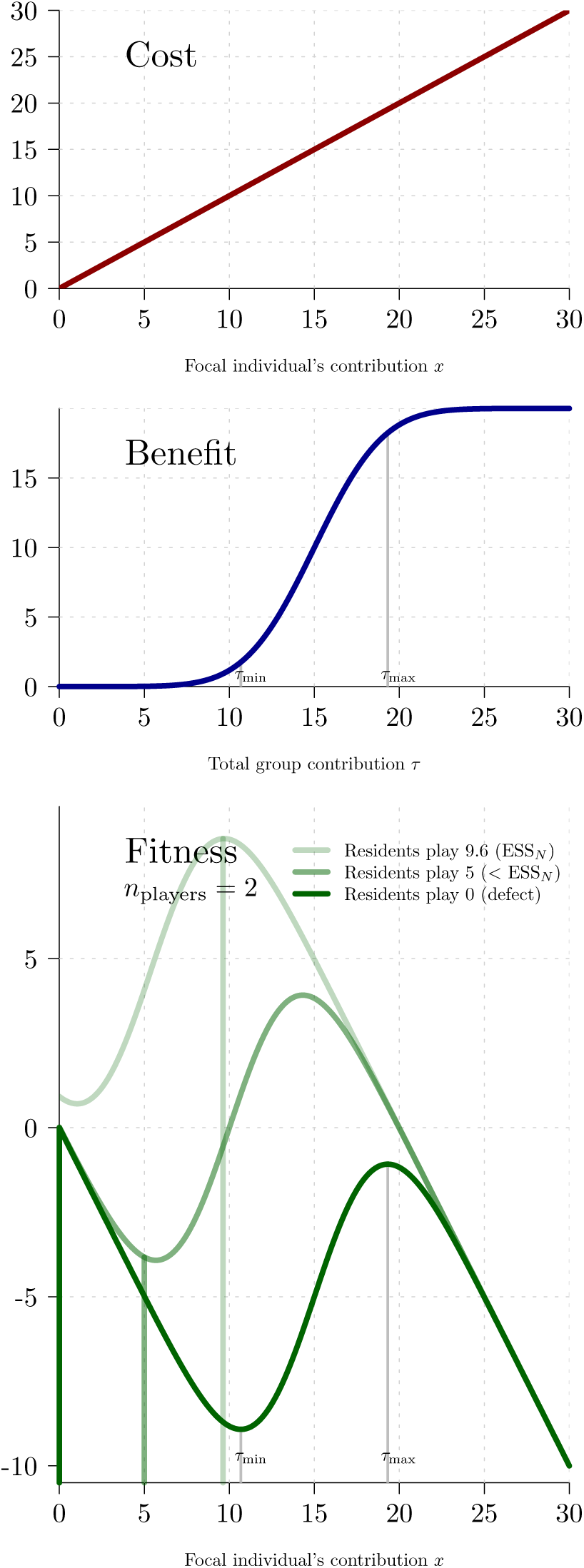
Example cost, benefit and fitness functions for a natural snowdrift game (NSG, defined in *Methods* §5.1). *Top panel:* The cost function is simply *C*(*x*) = *x. Middle panel:* The benefit function *B*(*τ*) is given in *Methods* equation (23); parameter values are *L* = 10, *k* = 1, *m* = 1.5, *τ*_turn_ = 15. *Bottom panel:* Fitness is shown for three situations involving groups of *n* = 2 individuals. (i) Residents cooperate and contribute the ESS_N_ (light green, *X*_res_ = 9.63) (ii) Residents cooperate but contribute less than the ESS_N_ (medium green, *X*_res_ = 5). (iii) Residents defect, *i.e.*, contribute nothing (dark green, *X*_res_ = 0). Resident strategies are indicated by vertical lines in the same colour as the associated fitness function. In the case of defecting residents, a focal individual’s fitness function does not depend on the group size (*n*) and has a local maximum at the maximizing total good (*τ*_max_ = 19.3, thin grey vertical line).

Traditionally, an ***evolutionarily stable strategy (ESS)*** is one such that, when adopted by the entire population, a single mutant individual playing a different strategy cannot invade the population (33). Because the phenotypic changes caused by mutations are often small, local ESSs are of particular interest: a population of individuals playing a ***local ESS*** is resistant to the invasion of a single individual playing a slightly different strategy. A strategy is ***convergently stable*** if a population playing a different strategy evolves toward it (34); convergence can be either global or local.

In infinite populations, the theory of adaptive dynamics (2, 8, 35) identifies a ***singular strategy*** as one at which the selection gradient, *∂*_*x*_*W (x, x* + (*n* - 1)*X)*|_*x*=*X*_, vanishes (36, Table 1); for an NSG, this reduces to

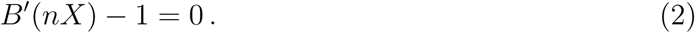

A singular strategy for which the mutant fitness is concave near the singular strategy is a local ESS.^*∗*^ Local convergent stability of singular strategies is also defined via a condition on the local fitness difference [see Table 1 of (2)].

The definition of singular strategies can be extended to finite populations: The defining feature of a singular strategy is that when it is played by a resident population, directional selection vanishes; for an NSG, this condition reduces to

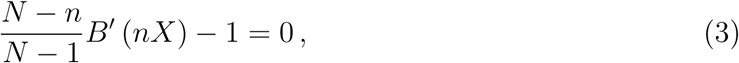

[see definition 4.3.5 of (37) and equation (28)]. The finite-population extension of the concept of evolutionary stability is more involved, because it must account for the fact that selection can favour fixation of a mutant strategy, even if selection opposes its invasion (9). Thus, the standard definition of an evolutionary stable strategy in a finite population (***ESS***_***N***_ (9)) requires that selection oppose both invasion by, and fixation of, mutant strategies. In addition, the presence of one or more mutants in a finite population has a non-negligible effect on the fitness of residents (whereas finitely many mutants cannot affect the mean fitness of residents in an infinite population).

Fixation probabilities depend on the ***selection process*** (31), *i.e.*, the stochastic process by which differences in fitnesses of individuals playing different strategies generate changes in the frequencies of strategies in the population over time. As a result, the strategies that are evolutionarily stable in finite populations depend on the selection process. Variants of the Moran and Wright-Fisher processes (23, 38, 39) are commonly assumed, but are idealizations that do not exactly describe realistic populations (*e.g.*, (40)). We are spared this complication in this paper because, for the games we consider, every ESS is a ***universal ESS***, that is, all ESSs are evolutionarily stable irrespective of the selection process. Consequently, we need not specify the population-genetic processes underlying selection, and we obtain general results about evolutionary stability. We use the term ***universal*** more generally to indicate that a property or statement holds for any selection process.

## 3 Results

### ESSs in infinite populations

As we have previously shown (30), if an NSG (*Methods* §5.1) is played in an infinite population then there are always two (and only two) ESSs:

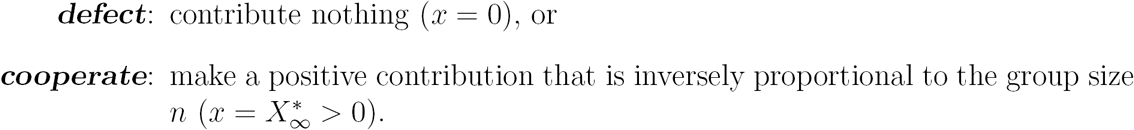

Both ESSs are global, and both are locally convergently stable [theorem 4.1 of (30)]. At the cooperative ESS, everyone contributes an equal share of the amount that maximizes individual fitness given that everyone contributes equally. In terms of this ***maximizing total good*** *τ*_max_ (see *Methods* §5.1 and figure 1), the cooperative ESS is

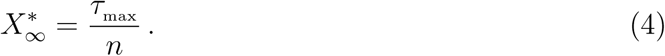

### ESSs in finite populations

In a finite population, NSGs do not necessarily have a cooperative ESS_N_, and when they do it is not necessarily possible to find an explicit formula for evolutionarily stable cooperation levels in terms of the parameters of an NSG (nevertheless, cooperative ESS_N_s are always easy to find numerically within the interval (6) identified in the following theorem).

#### Theorem 1

(Existence and universality of stable cooperation levels in the natural snowdrift game). *Consider a finite population (of N individuals) that is subject to selection resulting from groups of n individuals playing an NSG* [*defined in Methods §5.1*]. *A strategy X is singular if and only if*

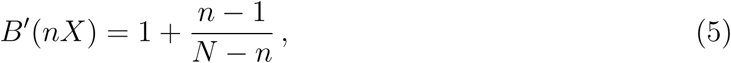

*and any such strategy X lies in the open interval*

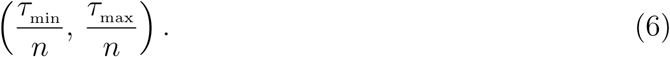

### Necessary condition for ESS_N_

*Any cooperative ESS*_*N*_ *(X >* 0*) satisfies both equation* (5) *(which implies* 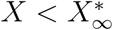 *) and*

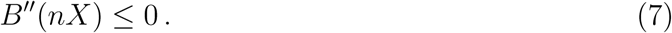

### Sufficient condition for universal ESS_N_

*If X satisfies equation* (5) *and*

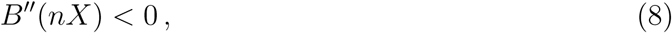

*then X is a universal ESS*_*N*_ *that is (universally) locally convergently stable.*

### ESS_N_s in large populations

*If B”* (*τ*_max_) ≠ 0 *and the group size n is either fixed, or satisfies 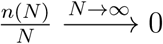, then for any sufficiently large population size N*, *there is a universal* 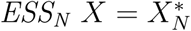 *satisfying inequality (8). Moreover*, 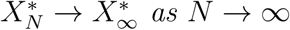.

While the evolutionarily stable cooperation levels in finite and infinite populations are never exactly the same, theorem 1 shows that the difference is negligible in sufficiently large populations if as the population size *N → ∞*, groups become a vanishingly small proportion of the population (*cf.* figure 2). However, if group size is not sufficiently small relative to the total population size then evolutionary predictions from finite population models differ qualitatively from the predictions for infinite ones: it may actually be impossible for cooperation to evolve at all. This is formalized in the next theorem.

**Figure 2:**
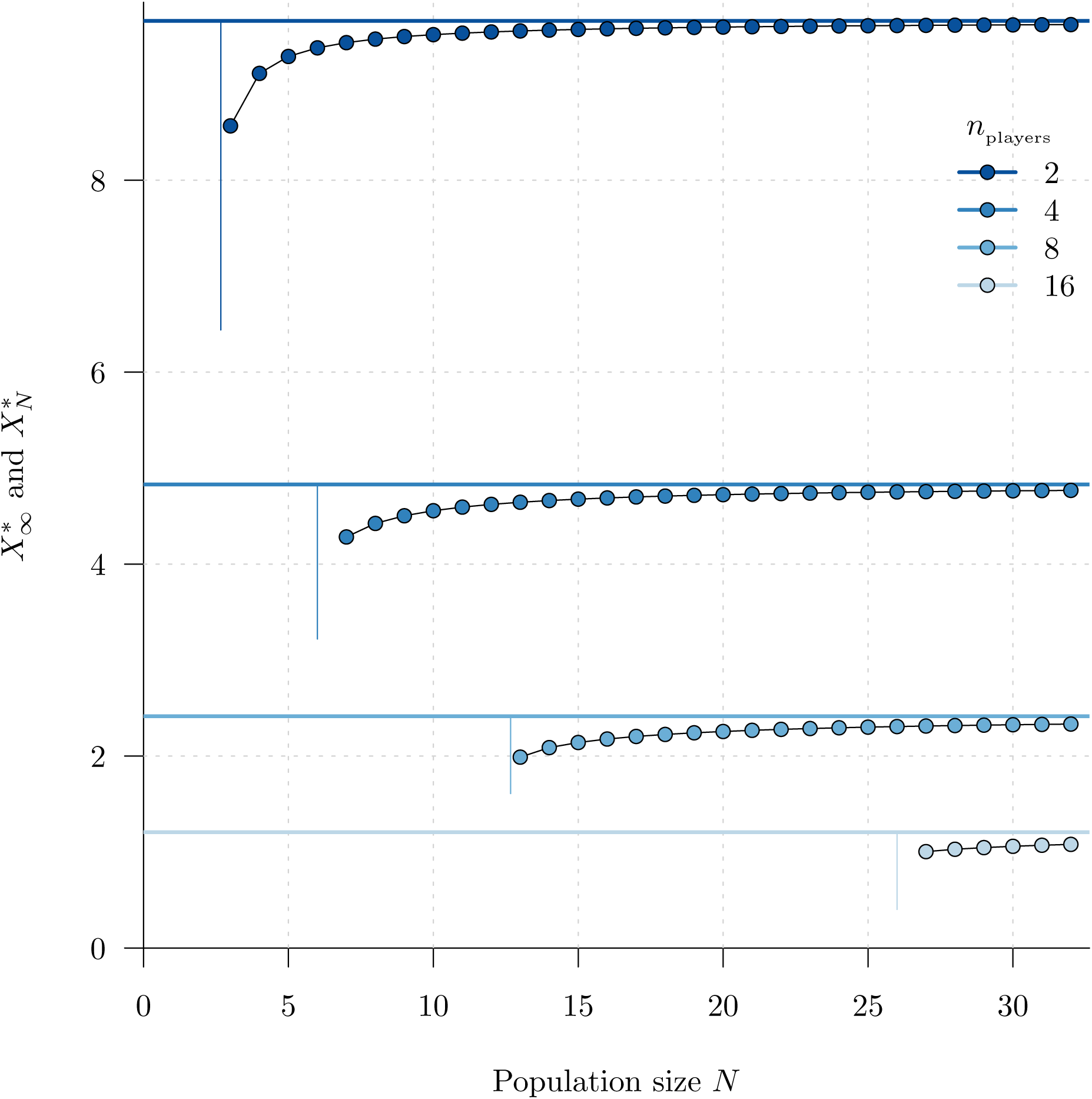
Evolutionarily stable strategies in the natural snowdrift game (*Methods* §5.1, with the sigmoidal benefit function shown in figure 1). For several group sizes (*n*), the infinite population ESS (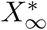, equation (4)) is shown as a horizontal line, and the finite population ESS_N_ (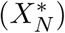) is shown with dots as a function of population size *N*. The vertical line segments indicate the critical population size threshold (*N*_min_, inequality (13)). A cooperative ESS_N_ exists if and only if *N > N*_min_.

#### Theorem 2

(ESS_N_s of the natural snowdrift game). *Consider a finite population (of N individuals) that is subject to selection resulting from groups of n individuals playing an NSG* [*defined in Methods §5.1 with fitness W defined by equation* (20)]. *Let m denote the* ***maximal marginal fitness***, i.e.,

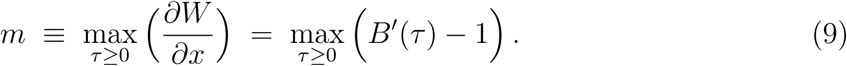

*Then m >* 0 *and there is a* ***critical maximal marginal fitness*** *threshold*,

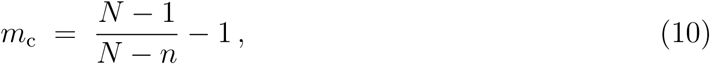

*such that*^*†*^

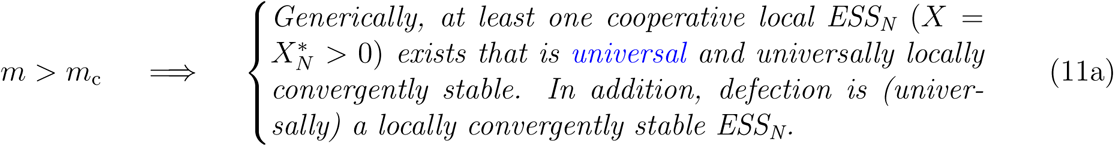

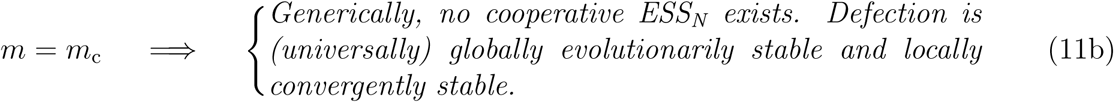

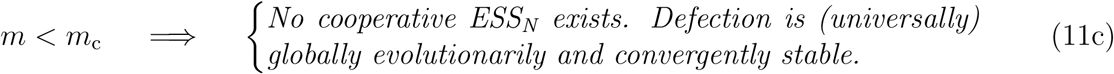

This theorem predicts qualitatively different evolutionary outcomes, depending on the maximal marginal fitness (*m*): Equation (10) gives the critical maximal marginal fitness above which a cooperative ESS_N_ exists, and below which defection is the only ESS_N_. Theorem 2 thus connects the maximal marginal fitness—a property of the fitness function that relates investments in the communal task to fitness benefits—with properties of the population of interacting agents: the population size (*N*), the number of players in a group (*n*), and the number of groups (*G* = *N/n*).

Equation (10) expresses the critical maximal marginal fitness in terms of a given population size and given group size. To clarify the roles of group size and number of groups in the evolution of cooperation, it is useful to think instead of the maximal marginal fitness (*m*) as given (*i.e.*, as a fixed property of the strategic interaction). Then, in the inequality *m > m*_c_ [see (11a)], we can replace *m*_c_ by the expression on the right hand size of Equation (10), and solve for a critical number of groups (*G*_c_) or critical group size (*n*_c_).

### ESS conditions in relation to the number of groups (*G*)

Condition (11a) can be expressed equivalently as

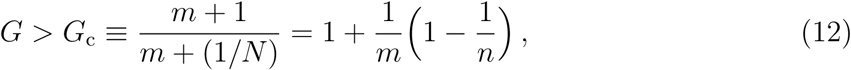

*i.e.*, the number of groups *G* must be greater than *G*_c_, the minimum number of groups that support cooperation in a population of size *N* (or in groups of *n* players). For any given number of players in a group (*n*), if we multiply inequality (12) by *n* we see that cooperation cannot evolve—*i.e.*, no cooperative ESS_N_ exists—*unless* the population size is *greater than* a critical population size^*‡*^,

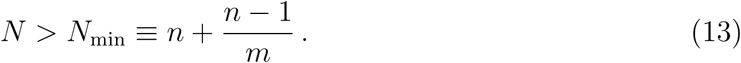

Figure 2 illustrates this result for the particular NSG specified by the benefit function shown in figure 1. Put another way, for a given group size *n*, if the population size *N* is too small then there is no cooperative ESS_N_, but if *N* is sufficiently large then there is a (universal) cooperative ESS_N_. For any given population size *N*, there are group sizes *n* and benefit functions *B*(*τ*) that yield *N*_min_ *> N*, so a qualitative difference between the evolutionary outcomes in finite and infinite populations can occur either for small or large population sizes.

### ESS conditions in relation to group size (*n*)

Rearranging condition (11a) again, we can write

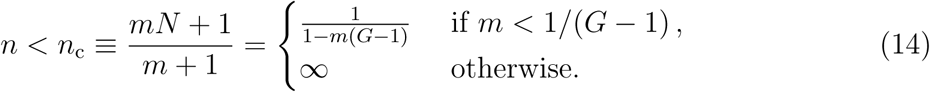

*i.e.*, for cooperation to evolve, the group size *n* must be less than *n*_c_, the maximum size of groups that support cooperation in a population of size *N* (or a population divided into *G* groups^*§*^). Multiplying inequality (14) by *G* and rearranging, we obtain

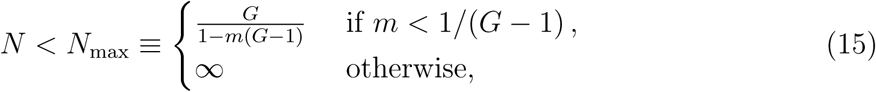

*i.e.*, if the number of groups is fixed (and smaller than 1+1*/m*) then in order for a cooperative ESS_N_ to exist, the population size must be *less than* the threshold in inequality (15), as illustrated in figure 3.

**Figure 3:**
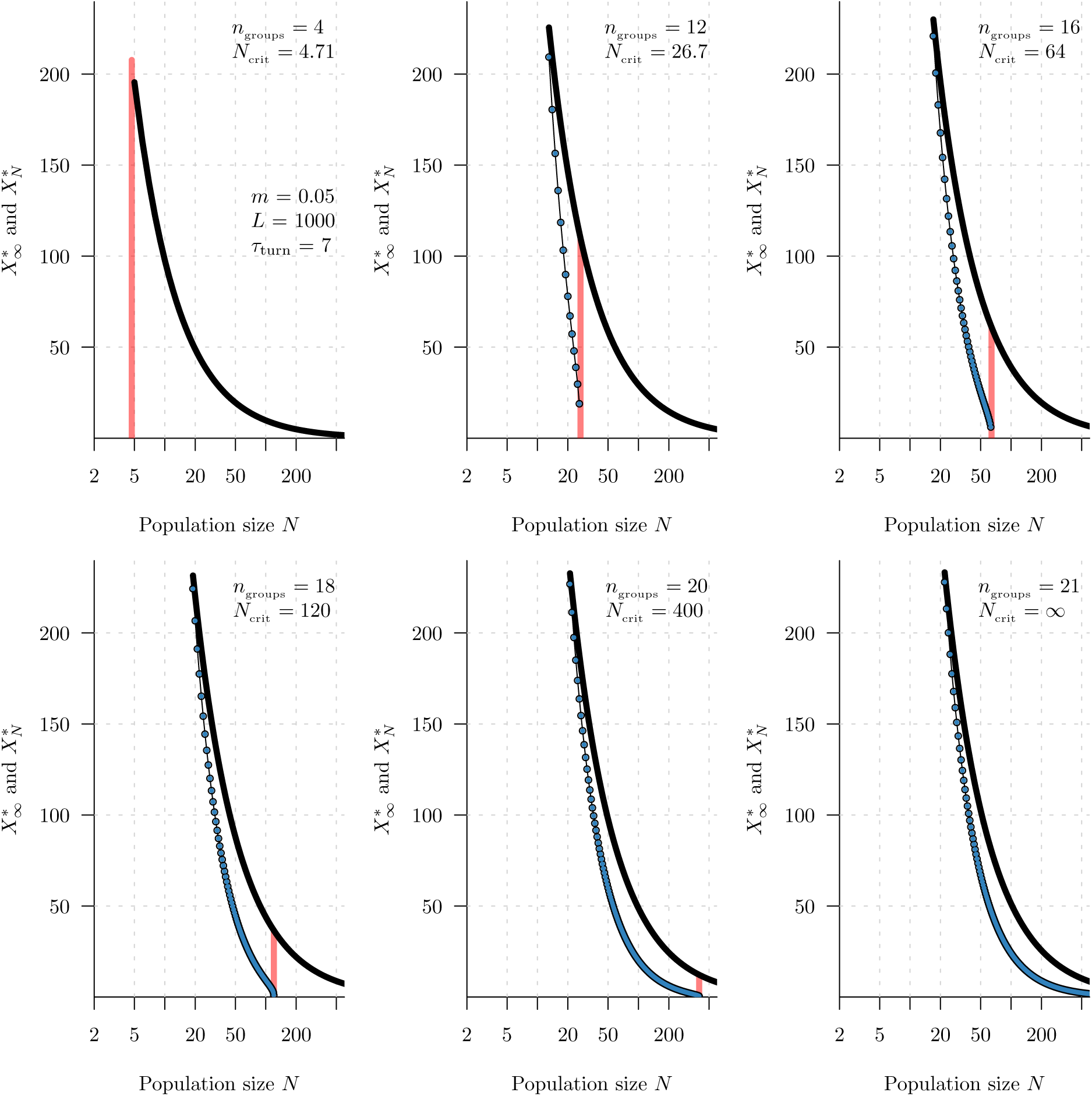
Evolutionarily stable strategies in the natural snowdrift game (*Methods* §5.1), with the sigmoidal benefit function *B*(*τ*) given in *Methods* equation (23); parameter values are *L* = 1000, *k* = 1, *m* = 0.05, *τ*_turn_ = 7. For several numbers of groups (*G*), the infinite population ESS (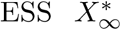, equation (4)) is shown as a curve, and the finite population ESS_N_ (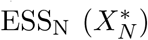 is shown with dots as a function of population size *N*. For each number of groups, the minimum population considered is *N* = *G* + 1. The vertical line segments indicate the critical population size threshold (*N*_max_, inequality (15)), below which a cooperative ESS_N_ exists (in contrast to the situation in which the group size *n* is fixed and an ESS_N_ exists only above a critical population size; *cf.* figure 2).

### Lack of ESS_N_ for any population size

It is even possible that there is a cooperative ESS if the population is infinite, but no cooperative ESS_N_ for any finite population size *N*. This is easy to verify for an NSG as follows. As noted above, an NSG always has an infinite-population cooperative ESS (4). An ESS_N_ exists if and only if inequality (11a) [or inequality (14) or inequality (12)] is satisfied. Rearranging inequality (12) [or equation (10)], we can write, equivalently,

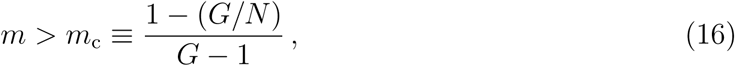

*i.e.*, there is a cooperative ESS_N_ if and only if the maximum marginal fitness *m* exceeds the threshold *m*_c_ (exactly the same threshold that appears in equation (10), but expressed here in terms of *G* rather than *n*). Suppose now that the population is divided into a given number of groups, *G ≥* 2. There must be at least two individuals in each group, so *N ≥* 2*G* and hence *G/N ≤* 1*/*2. Consequently, for *any possible population size N*, we have

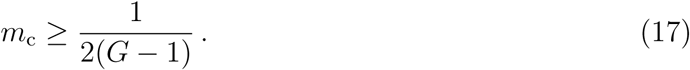

Therefore, if the benefit function is such that

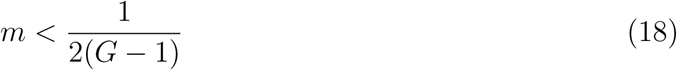

then no cooperative ESS_N_ exists, no matter how large the population size *N*. Yet, when the game defined by the same cost and benefit functions is played in an *infinite* population, a cooperative ESS exists (regardless of the group size *n*). Given *G*, in the example of the NSG defined using equation (23), it is easy to satisfy inequality (18) since the only constraint on *m* is that it must be positive.

Above, we have considered populations divided into a given number of groups. Alternatively, we could consider groups of a given size (*n*), and ask whether it is possible for a public goods game to have a cooperative ESS if the population is infinite but no cooperative ESS_N_ for any finite population size. As we show elsewhere, NSGs do not have this property, but there are snowdrift games that *do* have it (41).

### Confirmation with both selection and mutation

Lastly, in figure 4 we complement our rigorous analyses with individual-based simulations of finite populations in which individuals undergo both selection and mutation (see appendix 5.2 for details). Simulations such as these confirm that rigorous game-theoretical analyses—which are based on selection acting with only two types in the population—correctly predict evolutionary outcomes in realistic populations in which each individual can, in principle, be playing a different strategy.

**Figure 4:**
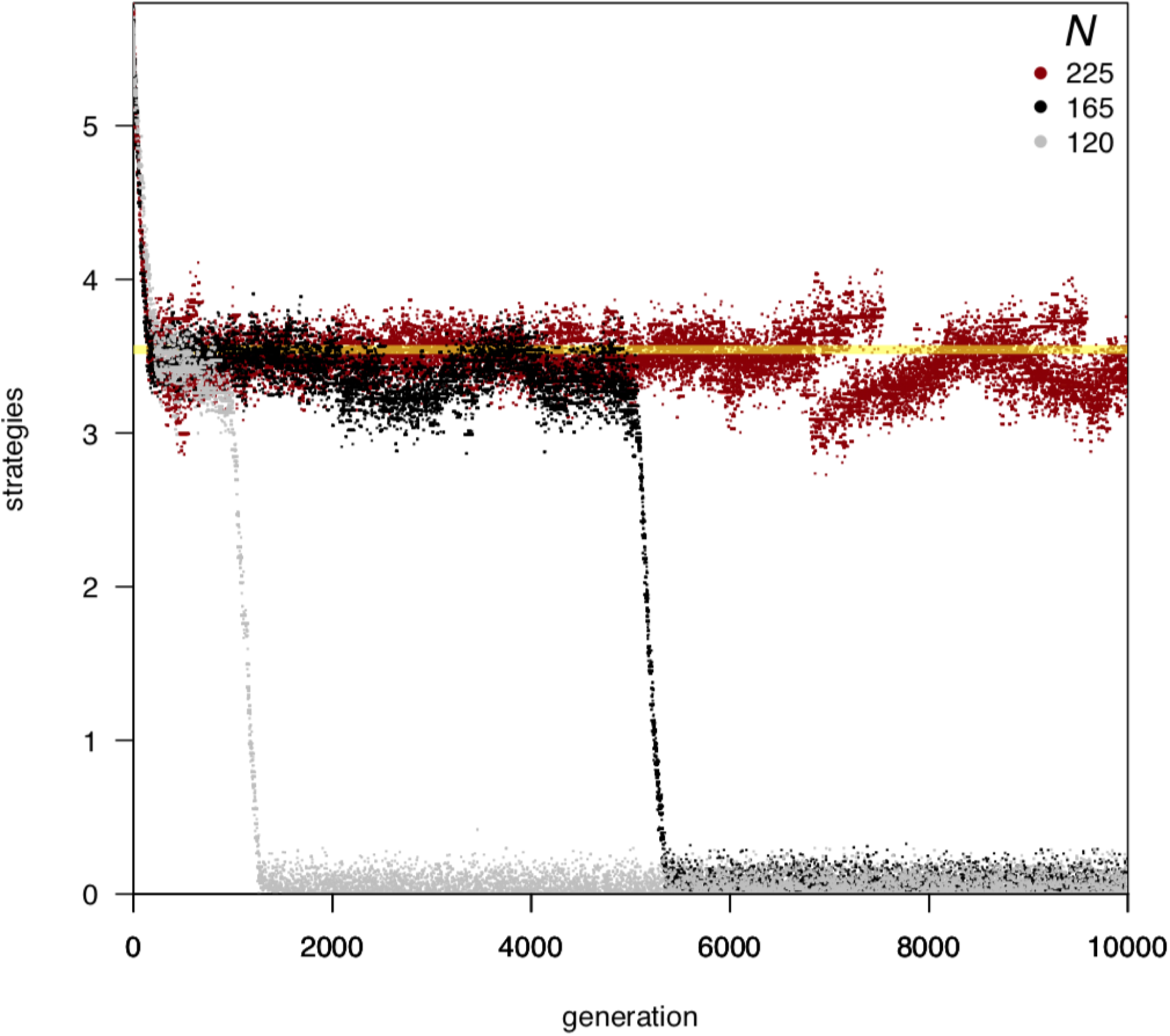
Individual-based simulations (details in appendix 5.2) of populations playing an NSG with cost and benefit functions as in figure 1 and group size *n* = 15, for population sizes *N* = 225 (red), 165 (black) and 120 (grey). The horizontal axis is the number of generations elapsed, and the vertical axis is the strategy (contribution level) of each individual in the population. The strategies present in the population in each generation are plotted on a vertical line intersecting the horizontal axis at the corresponding point. For *N* = 120, defecting is the unique, globally convergently stable ESS_N_; for *N >* 155, a cooperative ESS_N_ is predicted at *X*^*∗*^ = 3.54 (marked with a horizontal yellow line). The ESS for an infinite population playing this game is 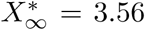. Note in these simulations, the mutation rate is high enough that populations contain more than two strategies at any given generation (in contrast to our rigorous mathematical analysis of dimorphic populations).

## 4 Discussion

We have seen that the evolutionary dynamics of the class of natural snowdrift games (NSGs, defined in *Methods* §5.1) are different when played in finite *vs.* infinite populations. Since all real populations are finite, it is important to understand how inferences based on infinite-population analyses of the multi-player snowdrift game (*e.g.*, (3, 30, 42)) might be affected. More generally, under what circumstances are infinite-population analyses of the evolution of cooperation likely to lead to invalid inferences about real populations?

We have shown that there are games for which it is *possible* that cooperation can evolve in an infinite population but not in any finite population (no matter how large). This extreme possibility emphasizes that inferences drawn from infinite population analyses should always be regarded cautiously.

The infinite-population approximation is *likely* to predict incorrect evolutionary outcomes if the number of individuals playing the game (the group size, *n*) is substantial relative to the total population size (*N*). Exactly what “substantial” means will depend on the game in question and the population size; we have specified this threshold precisely for NSGs in inequality (14). Evolutionary predictions derived from infinite population analyses can be incorrect for finite populations of any size (figure 2 and theorem 2). The origin of such erroneous inferences is that finite groups (no matter how large) are always negligible in size compared to an infinite underlying population, but not compared to a finite underlying population.

Intuition for how different predictions arise in finite and infinite populations can be developed by considering a thought experiment in which the population (of size *N*) is *simultaneously* divided into *G* groups that play the game. If a single mutant invades the resident population, the probability that a randomly chosen group contains the mutant is 1*/G*. If the population size were then increased by adding more and more groups of the same size (*G→ ∞*, keeping *n* fixed), then the effect of the mutant on the residents would be “infinitely diluted” (the mutant would have a negligible effect on residents’ fitnesses as *N→ ∞*). If, instead, the population size were increased by adding individuals to the existing groups (without increasing the number of groups) then the probability that a randomly selected group contains the mutant would not change; however, in this version of the thought experiment, the limit *N→ ∞* entails the size of each group also becoming infinitely large.

Adaptive dynamics, which has been extensively used in the study of evolutionary dynamics [*e.g.*, (3, 42, 43), as well as (44) and references therein], relies on an infinite-population approximation (8). Previous work has presented reasonable arguments to justify this approximation (*e.g.*, (35)) and reported general agreement between adaptive dynamics and stochastic simulations of finite populations (see (45) for a review). In addition, specific agreement has been noted (15) between the finite-and infinite-population evolutionary dynamics of the multi-player snowdrift game with *discrete* strategies. These results appear to contrast those presented here, though (15) did observe that defectors prevail when the group size approaches the population size (even in situations in which cooperators and defectors can coexist in an infinite population). In other work, there has been a focus on situations in which the group size is much smaller than the population size, which reduces the chance of discovering discrepancies between finite and infinite population evolutionary predictions.

Our analysis of the class of natural snowdrift games is rigorous (theorems 1 and 2), and our conditions for existence of a cooperative ESS_N_ are universal (in the sense of being entirely independent of the selection process). Broadly, our results indicate that approximating large populations by infinite ones may generate misleading conclusions. In particular, inferences based on adaptive dynamics are not necessarily applicable to real (finite) populations. There is a general need to reevaluate the theoretical justification for approximating large populations by infinite ones, and to derive clear conditions for when such approximations are useful.

## 5 Methods

### 5.1 The natural snowdrift game (NSG)

This biologically motivated version of the continuous snowdrift game (§2) was introduced in (30). We consider a population of individuals that are identical except (possibly) with respect to the strategy (contribution level) adopted when playing the snowdrift game. In particular, there is no age, spatial, social or other structure in the population. Evolution affects only the contribution levels of individuals, so at any time the population is completely characterized by the set of strategies present in the population and the numbers of individuals (or population proportions) playing each strategy. An individual’s fitness is determined entirely by its payoff from the continuous snowdrift game played in groups of *n* individuals. We say that this population plays a ***natural snowdrift game*** (NSG) if, in addition, the cost and benefit functions have the following properties (which are satisfied by the example shown in figure 1):

a. The cost to the focal individual of a contribution *x* is measured in units of its impact on this individual’s fitness, that is,

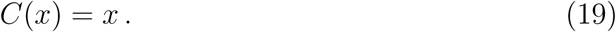

Thus, the focal individual’s fitness is

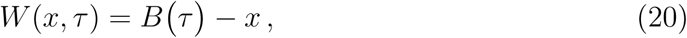

where *τ* is the total contribution in the focal individual’s group.
b. The benefit *B*(*τ*) is a *smooth* function of the total contribution *τ* (more precisely, *B* (*τ*) exists for all *τ ≥* 0).
c. There exist total contribution levels *τ*_min_ and *τ*_max_ (0 *≤ τ*_min_ *< τ*_max_) such that *B*(*τ*) *-τ* decreases for *τ < τ*_min_ and *τ > τ*_max_ and increases for *τ*_min_ *< τ < τ*_max_. Consequently, given condition (a), if only one member of a group contributes anything then that individual’s fitness [take *x* = *τ* in equation (20)] is locally minimized (maximized) if its contribution is *x* = *τ*_min_ (*τ*_max_).
d. There is a net fitness cost to an an individual who contributes *τ*_max_ when all other group members contribute nothing,

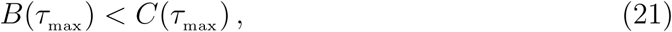

but there is a net incremental fitness benefit for contributing *τ*_max_ */n* if other group members contribute that amount,

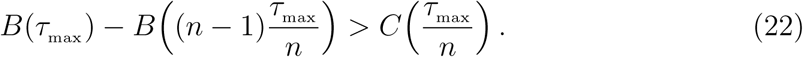

In an infinite population, condition (c) implies that *τ*_max_ */n* and 0 are the only local ESSs (30). Adding condition (d) guarantees that they are both global ESSs [0 via and condition (21) and *τ*_max_ */n* via condition (22); see (30)].

#### 5.1.1 Benefit function used for numerical examples

For the purpose of making example graphs and running simulations, we have used sigmoidal benefit functions. The biological motivation for this is that one would expect a nonlinear increase in the ease of passing the barrier as more snow is cleared, but eventually there can be no further benefit from additional work because all the snow has been cleared.

Specifically, for any integer *k >* 0 and real numbers *m >* 0, *L >* 0 and *τ*_turn_ *≥* 0, consider the benefit function

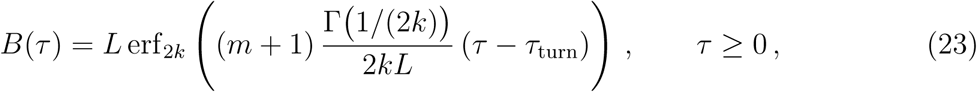

where erf_ℓ_ (*x*) is the generalized error function (46) of order ℓ,

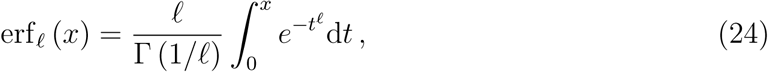

and Γ(*x*) is the gamma function [equation (50a)]. We analyze this flexible class of sigmoidal benefit functions in appendix B, where we show that the parameters *L* and *τ*_turn_ are the horizontal asymptote and the inflection point, respectively, *k* controls the “width” of the sigmoid^¶^, and *m* + 1 is the maximal marginal benefit (so that *m* is the maximal marginal fitness that results from this functional form, justifying our notation).

Figure 1 shows the benefit function (23) for particular values of *k, m, L* and *τ*_turn_, together with the corresponding fitness function (20) that results if residents defect, or—in groups of two individuals—if residents play the infinite population ESS [equation (4)]. Based on equation (23), in appendix B we derive explicit formulae for *τ* _min_ *τ*_max_, and 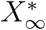 and 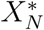 (interms of *m, L, τ*_turn_ and *k*).

The class of sigmoids based on generalized error functions is much more flexible than the more common “logistic” sigmoid used by (30, 42) (which is based on shifting, and horizontally and vertically stretching, the hyperbolic tangent function, tanh(*x*)). Whereas the maximum slope, horizontal asymptote and position of the inflection point uniquely determine the “width” of a logistic sigmoid, the generalized error function allows the width to be set independently via the parameter *k* [see equation (60)].

### 5.2 Individual-based simulations

The three individual-based simulations shown in figure 4 [for population sizes *N* = 120 (grey), 165 (black) and 250 (red)] were run using algorithm 1, which we implemented in an R (47) package. In the following description, we denote the normal distribution truncated to the interval (*l, u*) by `TruncNormal`(*µ, σ, l, u*). It is a assumed that values of the following parameters have been set:

- Parameters (*k, m, L* and *τ*_turn_) of the benefit function (23).
- Group size (*n*) and population size (*N*), such that *G* = *N/n* is an integer.
- Number of repetitions of the NSG between reproductive events (*n*_reps_).
- Maximum number of generations to evolve (*nGen*).
- Upper bound for contribution level (*x*_max_).
- Mean (*µ*_*x*_) and standard deviation (*σ*_*x*_) of an underlying Normal(*µ*_*x*_, *σ*_*x*_) distribution of strategies; the initial strategies (*x*_*i*_, *i* = 1, *…*, *N*) are to be sampled from `TruncNormal`(*µ*_*x*_, *σ*_*x*_, 0, *x*_max_).
- Mutation probability (*p*_mut_) per individual per generation.
- Standard deviation (*σ*) of an underlying Normal(0, *σ*) distribution of the strategy changes caused by mutations, and upper and lower bounds on mutation sizes, (*l, u*); when an individual playing strategy *x* mutates, its new strategy is sampled from `TruncNormal`(*x, σ*, max {0, *x - l*}, min {*x*_max_, *x* + *u*}), so that the mutation is within the interval [*l, u*] and the mutated strategy is in [0, *x*_max_].

## Acknowledgments

DE was supported by the Natural Sciences and Engineering Research Council of Canada (NSERC). CM was supported by the Ontario Trillium Foundation, the United States Defense Advanced Research Project Agency NGS2 program (grant no. D17AC00005), and the Army Research Office (grant no. W911NF1810325). We are grateful to Sigal Balshine, Ben Bolker, Michael Doebeli, Jonathan Dushoff, Gil Henriques, Paul Higgs and Rufus Johnstone for valuable discussions and comments.

## Algorithm 1 Individual-based simulation algorithm

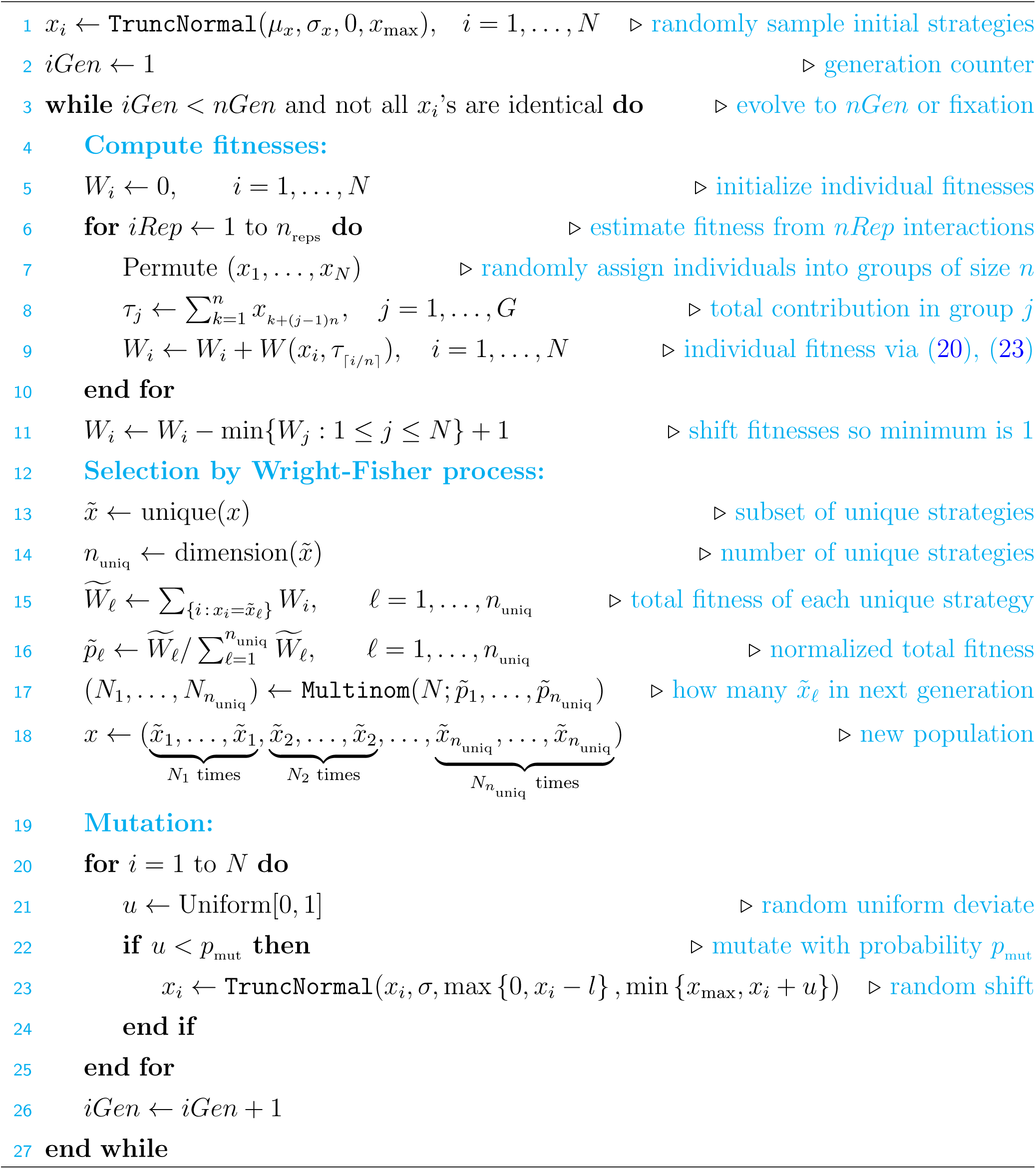

## SUPPORTING INFORMATION

### A Proofs

#### A.1 Analysis of the natural snowdrift game (NSG; § 5.1) in a finite population

Our main results are stated in theorems 1 and 2 (§ 3). Before developing the proofs in detail, it is useful to note that:

- *τ*_min_ *>* 0 (where *τ*_min_ is defined in assumption (c) of the definition of the NSG, § 5.1). To see this, suppose that *τ*_min_ = 0. Then assumption (c) implies that *B*(*τ*_max_) *≥τ*_max_, contradicting assumption (d).
- The benefit function *B*(*τ*) is twice-differentiable. This follows from assumption (b) in the definition of the NSG (§ 5.1).
- *B’* (*τ*_min_) = *B’* (*τ*_max_) = 1, *B’* (*τ*) *>* 1 for *τ*_min_ *< τ < τ*_max_, and *B’* (*τ*) *<* 1 otherwise [these properties of *B*(*τ*) follow from assumption (c)]. Consequently, *m >* 0 and *B”* (*τ*_max_) *≤* 0.

#### A.1.1 The mean fitness difference between mutants and residents

Consider a population of *N* individuals, comprised of *M*_p_ mutants who play *x* and *N - M*_p_ residents who play *X*, and denote the proportion of mutants in the population by ϵ = *M*_p_*/N*. Suppose that groups of *n* individuals are randomly sampled from this population without replacement, which implies that the number of mutants in each such group is hypergeometrically distributed with parameters *N*, *M*_p_ and *n* (37, 48); thus, the probability of *k* mutants occurring in a random sample of *n* individuals is

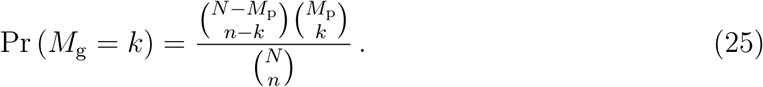

Suppose, moreover, that a focal individual is selected from the population by first sampling a group of *n* individuals, and then selecting one of the members of this group. Lastly, suppose for simplicity that individual fitnesses are given by the payoffs from a single round of the NSG played in such randomly selected groups^#^. We show elsewhere (37, eq. 4.61, p. 137) that the expected difference between the mutant and resident fitnesses is then

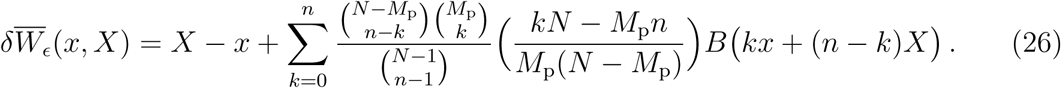

Differentiating equation (26) yields

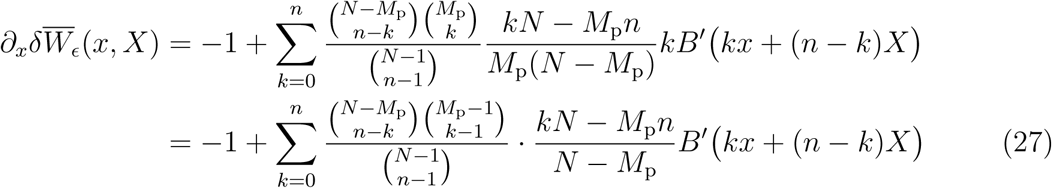

Differentiating with respect to *x* and setting *x* = *X*, we find (37, pp. 138–139)

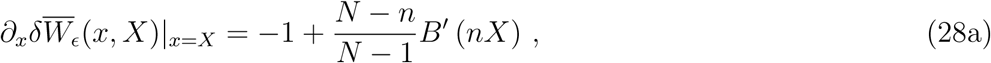

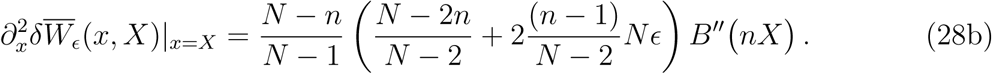

From these expressions we see that

- 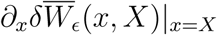 is independent of *ϵ*, and
- 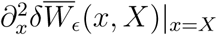 is linear in *ϵ*.

We will exploit these facts below.

#### A.1.2 Evolutionary and convergent stability of defection

##### Lemma 3

(Evolutionary stability of defection). *If the NSG (§ 5.1) is played in a finite population then not contributing (X* = 0*) is a locally convergently stable ESS*_*N*_ *for any selection process. Moreover, if the population and group sizes are the same (N* = *n, so the entire population plays the game together) then defecting is the unique ESS*_*N*_ *and is globally evolutionarily and convergently stable.*

*Proof. B’* (0) *<* 1 because *B*(*τ*) *- τ* decreases for 0 *≤ τ < τ*_min_, so using equation (28a),

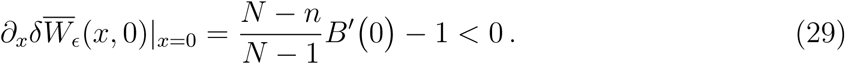

Because 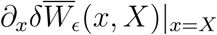 is continuous in *X*,

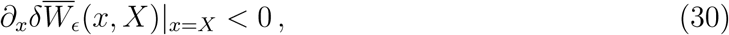

for *X* sufficiently small. From Theorem 4.3.9 in (37), it follows that *X* = 0 (defection) is convergently stable, and selection opposes invasion of mutants contributing a sufficiently small but positive amount, *x >* 0. To establish that *X* = 0 is evolutionarily stable, observe that equation (29) implies that 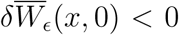 for sufficiently small *x*, so such mutants are selected against, regardless of their proportion (*ϵ*) in the population. Thus, corollary 5.4 of (31) implies that selection also opposes the fixation of such mutants.

Now suppose groups constitute the entire population, *i.e., N* = *n*. Then, for any resident strategy *X >* 0 and any number of mutants *M*_p_ *∈* {1, 2, *…*, *N -* 1}, mutants contributing less than residents to the public good (0 *≤ x < X*) have a higher payoff than residents; hence defection is the unique ESS_N_ and is globally convergently stable. Defection is also globally evolutionarily stable because for any mutant strategy *x >* 0 and any number of mutants (*M*_p_ *< N*), residents obtain a higher payoff than mutants (because they receive the same benefit without paying a cost).

#### A.1.3 Proof of theorem 1

Inserting equation (28a) into the definition of an evolutionarily singular strategy (definition 4.3.5 of (37)) implies that cooperative singular strategies are characterized by equation (5). Any solution of equation (5) must satisfy 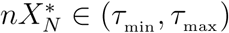, because the right hand side of equation (5) is greater than 1 and, as noted above, if *τ* ∉ (*τ*_min_, *τ*_max_) then *B’* (*τ*) *≤* 1.

##### Necessary condition for ESS_N_

Suppose that *X* solves equation (5) but *B”* (*nX*) *>* 0. Plugging equation (5) into equation (28a) gives 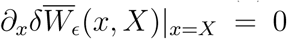. Rearranging equation (28b), we have

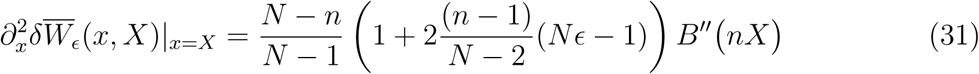

so 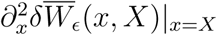 is increasing in *ϵ* and positive for any *ϵ ≥* 1*/N* (*i.e.*, any mixed population). Thus, when mutants play *x* sufficiently close to 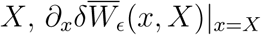 is negative for *x < X* and positive for *x > X*; hence, since 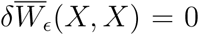, we must have 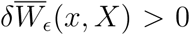 for any *x* that is near but not equal to *X* (and this is true for any number of mutants *M*_p_ = 1, *…*, *N -* 1). Corollary 5.4 of (31) then implies that selection favours the fixation of such mutants, so *X* is not an ESS_N_, regardless of the selection process. Thus, if 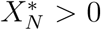 is an ESS_N_ then it cannot be that *B”* (*nX*) *>* 0, *i.e.*, inequality (7) holds.

##### Sufficient condition for universal ESS_N_

The sufficient condition for local universal evolutionary and convergent stability follows immediately from theorem 4.D.1 of (37) and equation (28).

##### ESS_N_s in large populations

Suppose that *B”* (*τ*_max_) ≠ 0 and consider the equation

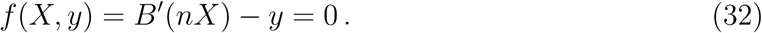

Noting that *f* (*τ*_max_ */n*, 1) = 0 and that

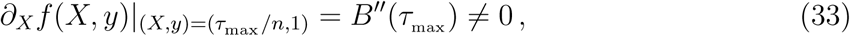

from the implicit function theorem (49, Theorem 12.40), there exists a differentiable function *X*(*y*) defined in a neighbourhood of *y* = 1, such that *X*(1) = *τ*_max_ */n* and

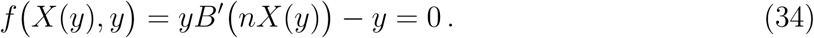

Now suppose that the group size *n* is either fixed, or varies with population size but satisfies

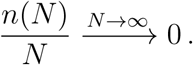

If we define 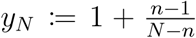 then *y*_*N*_ *→* 1, so for all sufficiently large population sizes *N*, equation (34) can be solved implicitly for 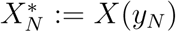. Such 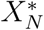 then solve equation (5), and 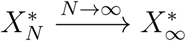 because *X*(*y*) is continuous. Recalling that *B”* (*τ*_max_) *≤* 0 and *B* ^*”*^ (*τ*_max_) ≠ 0 by assumption, we have *B”* (*τ*_max_) *<* 0, so for sufficiently large 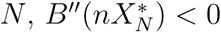. Theorem 4.D.1 of (37) then implies that for sufficiently large *N*, 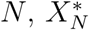 is a universal local ESS_N_ and is locally convergently stable.

□

#### A.1.4 Proof of theorem 2

First, note that *X* = 0 is always a locally convergently stable ESS_N_ (lemma 3). From Corollary 4.3.8 of (37), selection opposes invasion of a cooperative resident strategy *X >* 0 by sufficiently similar mutant strategies only if *X* is singular, which (using equation (28a)) occurs *iff X* satisfies

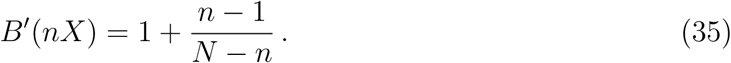

Because *B’* (*nX*) *>* 1 only if 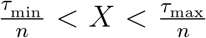, if a cooperative ESS_N_ exists then it must lie in this interval.

**Case *m > m***_**c**_. Because *B’* (*τ*_max_) = 1 and *B’* (*nX*) is a continuous function of *X* on the interval [*τ*_min_ */n, τ*_max_ */n*], it follows from the intermediate value theorem (49) that equation (5) has a solution in this interval. Let be the set of singular strategies, *i.e.*, solutions of equation (5),

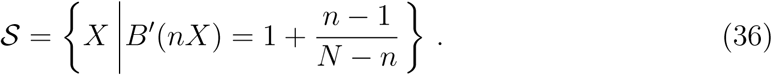

Note that from theorem 1, *S ⊂* (*τ*_min_ */n, τ*_max_ */n*). Denote the largest solution of equation (35) by 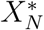, *i.e.*,

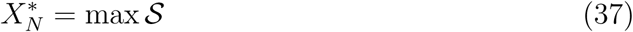

(this maximum exists because the continuity of *B*^*’*^ (*nX*) on a closed interval implies sup *S ∈ S*).

Generically ^‖^, 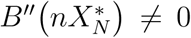. We claim that 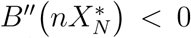. To see this, suppose, in order to derive a contradiction, that 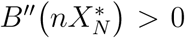. Then, *B’*(*nX*) increases in a neighbourhood of 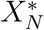, so there exists 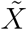 such that 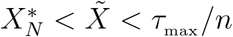 and

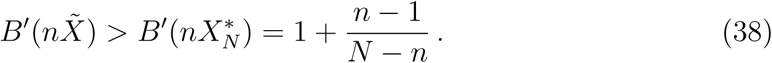

From the intermediate value theorem, there exists *X ∈ S*

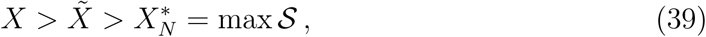

a contradiction.

Thus 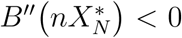 and *C”* (*X*) = 0, so theorems 4.D.1 and 4.3.9 of (37) imply that 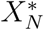 is a local ESS_N_ and is locally convergently stable.

**Case *m* = *m***_**c**_. Suppose, in order to derive a contradiction, that *X >* 0 is a cooperative ESS_N_. From theorem 1, *X* must solve equation (35) so, from the definition of *m*_c_ in equation (10),

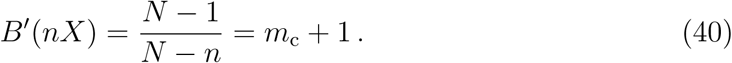

Suppose further that arg max *B’* (*τ*) does not contain an interval (*i.e.*, the marginal benefit *B’* is not maximal for an interval of total contributions *τ*), which happens generically. Then, any total contribution in arg max *B* ^*’*^ (*τ*) is a local maximum of *B’* (*τ*). It follows that if *x < X* and *x* is sufficiently close to *X*, then

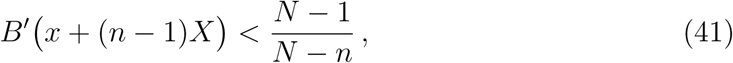

and therefore from equation (27),

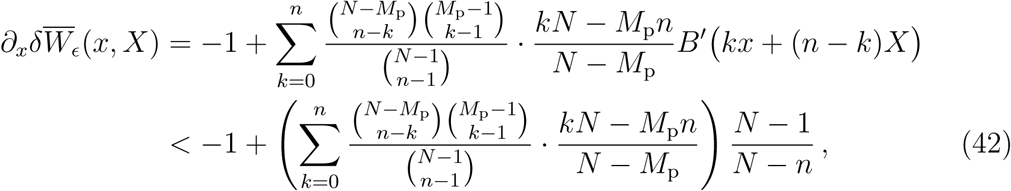

which, together with the identity (37, equation (4.63), p. 138),

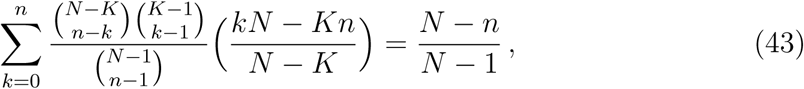

implies that 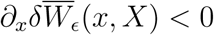. Hence, similar to an argument in the proof of theorem 1, since 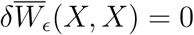, we must have 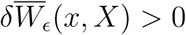 for any *x* that is slightly less than but not equal to *X* (and this is true for any number of mutants *M*_p_ = 1, *…*, *N –*1). Consequently, selection favours the invasion and replacement of *X* by any such *x*, so *X* is not evolutionarily stable.

To see that defection is globally evolutionarily stable, substitute *X* = 0 in equation (27) to get

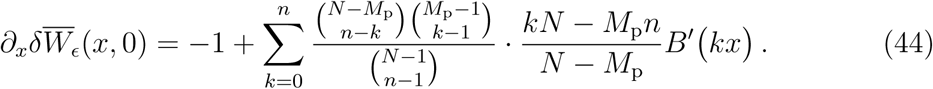

Noting that for all *x >* 0, *B kx ≤ m*_c_ + 1, we have

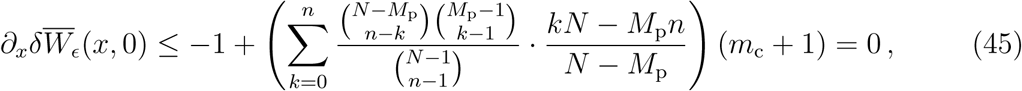

where we have used equations (10) and (43) in the last equality. Thus, 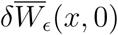 is non-decreasing in *x*. Moreover, if *x < τ*_min_ */n*, then *B’ kx <* 1 for all *k* = 0, *…*, *n*, so similarly, equations (43) and (44) imply that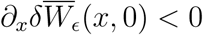. Because 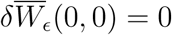, it follows that 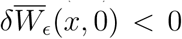 for all *x >* 0 (regardless of the proportion of mutants in the population). Thus, from (31, corollary 5.4), when residents defect, selection opposes invasion and fixation of any mutants.

**Case *m < m***_**c**_. In this case, equation (5) has no solution, and no cooperative ESS_N_ exists.

To see that defection (*X* = 0) is globally evolutionarily and convergently stable, observe first that *m < m*_c_ implies

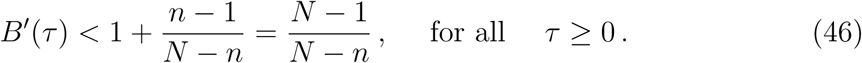

Then, using equations (27), (46) and equation (4.63) on p.138 of (37), it follows that

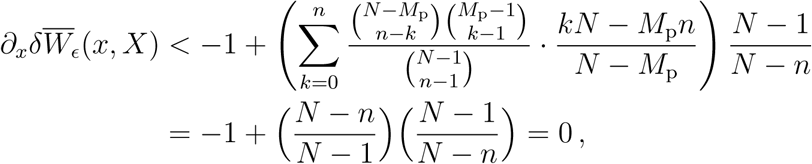

so 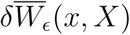 decreases with *x ≥*0 for any *X ≥*0. Thus, from (31, corollary 5.4), defection (*X* = 0) is a globally evolutionarily and convergently stable strategy.

□

### B Analysis of the benefit function used for numerical examples

In this appendix we define the class of sigmoidal benefit functions that we have used to illustrate our results, and derive a variety of analytical formulae that we have found useful when working with these functions.

#### B.1 Sigmoids using generalized error functions

For any integer *k >* 0 and real *m >* 0, *L >* 0 and *τ*_turn_ *≥* 0, consider the benefit function

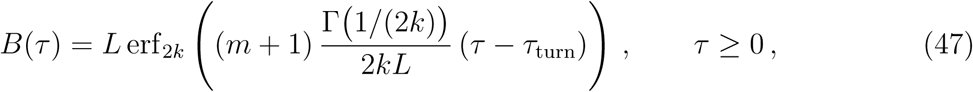

where erf_ℓ_ (*x*) is the generalized error function of order ℓ,

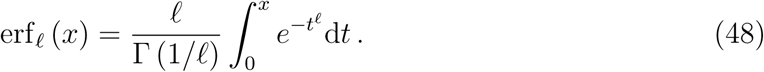

This class of functions generalizes the the error function, erf, which is recovered for ℓ = 2 or, equivalently, *k* = 1; see § B.2.

#### Expressing generalized error functions using gamma functions

It is sometimes convenient to express erf_ℓ_ in terms of gamma functions. For *x >* 0, the transformation *z* = *t*^ℓ^ (*t* = *z*^1/ℓ^ and 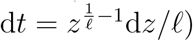 gives

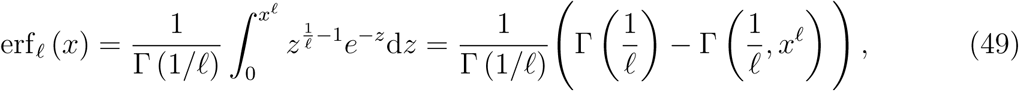

Where

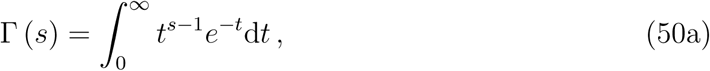

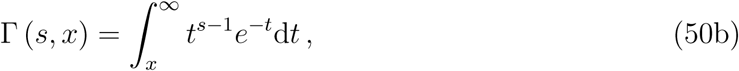

are the gamma^*∗∗*^, and upper incomplete gamma functions, respectively. Note that we are only interested in generalized error functions of even order (ℓ = 2*k*), which are odd functions of *x*.

#### Parameter meanings

Because equation (49) implies

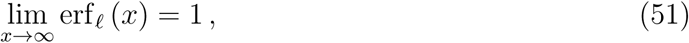

it follows that

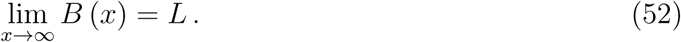

We show below that the inflection point of *B* (47) is *τ*_turn_, and that the maximal marginal fitness given the benefit function *B* is *m*.

From the integral definition of the generalized error function [equation (48)]

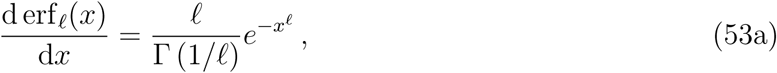

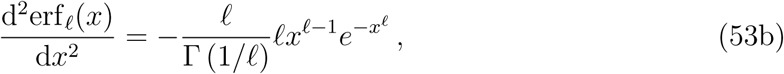

so

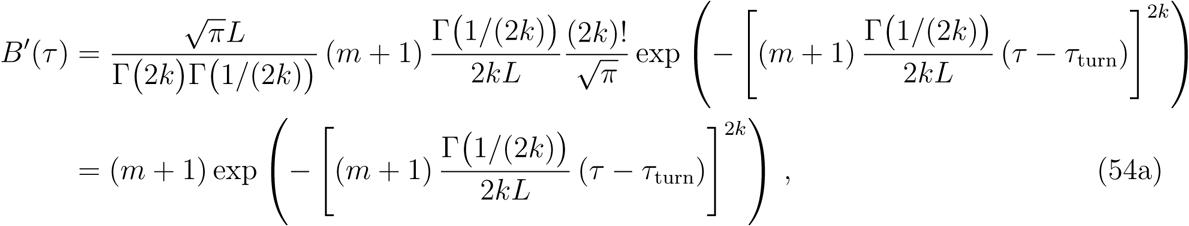

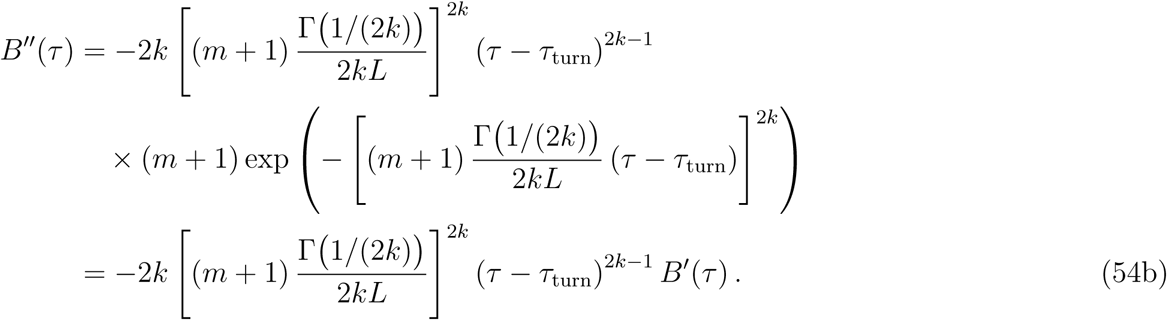

Consequently, *τ*_turn_ is the unique solution of *B*^*″*^ (*τ*) = 0, and is thus the only inflection point. *B’* (*τ*) is always positive, and hence *B* (*τ*) is monotonically increasing. However, *B″* (*τ*) *>* 0 for *τ < τ*_turn_ and *B″* (*τ*) *<* 0 for *τ > τ*_turn_, and hence

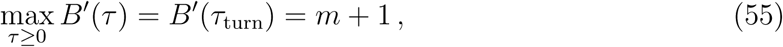

so from equation (9), the maximal marginal fitness is

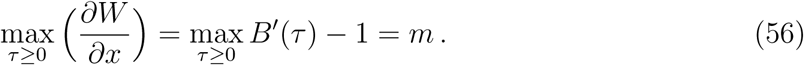

#### The minimizing and maximizing total goods

Since *B’* (*τ*) is monotonic on each of the intervals, (–*∞, τ*_turn_) and (*τ*_turn_, *∞*) and *B’* (*τ*) is even, for any *b ∈B’* (ℝ_*≥*0_) = (0, *m* + 1], we can find two real values of *τ* for which *B’* (*τ*) = *b* (although one of these values may be negative and therefore biologically irrelevant, because total contributions to the public good cannot be negative). To find these values of total contribution *τ*, we set *B’* (*τ*) = *b* in equation (54a), and get

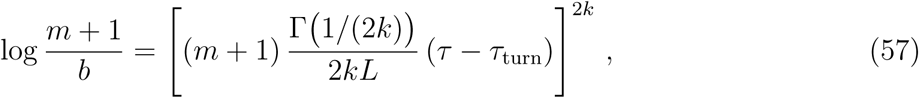

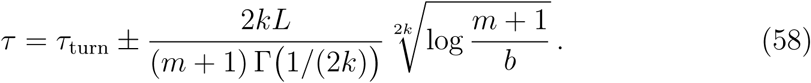

To find *τ*_max_ and *τ*_min_, we substitute *b* = *B* ^*’*^ (*τ*) = 1 in equation (58) and, noting that *B*^*″*^ (*τ*) changes sign from positive to negative at *τ*_turn_, we have

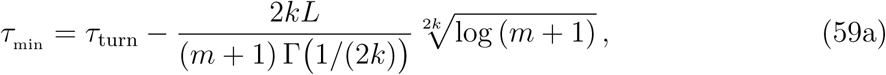

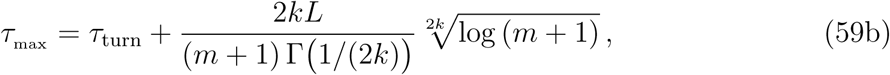

and the distance between the location of the fitness minimum and maximum is

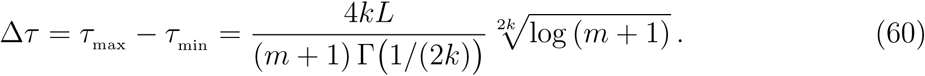

#### The infinite-population cooperative ESS

equation (4) then gives

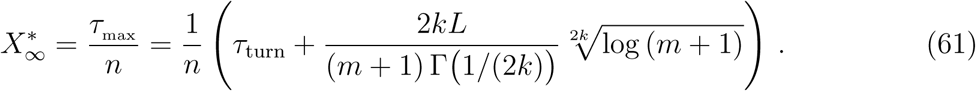

Using *B’* (*τ*_max_) = 1 and equation (59b) in equation (54b), we have

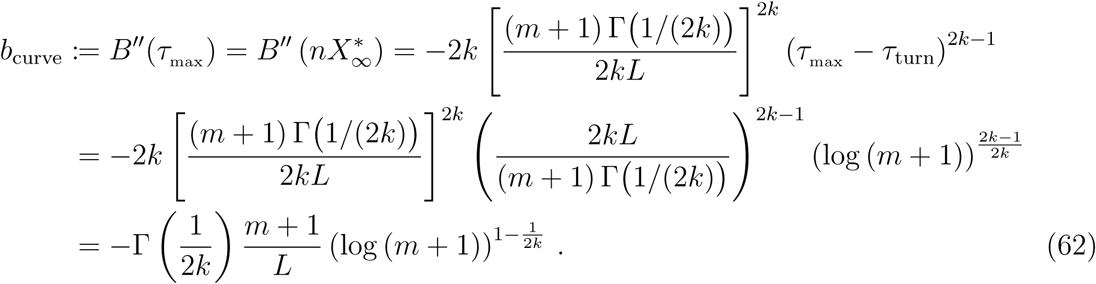

Using equation (47) and the fact that erf_2*k*_ is odd,

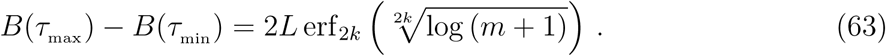

#### Singular and evolutionarily stable cooperative strategies in finite populations

In a finite population of size *N*, a singular strategy 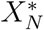 of the NSG is a solution of equation (5), that is,

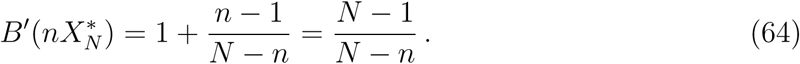

so equation (58) implies that at the ESS, the total contribution must be one of

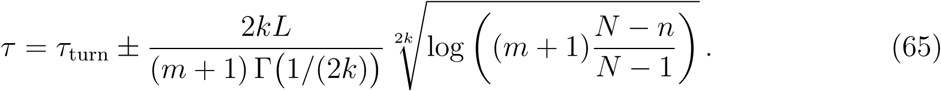

There are therefore two singular strategies,

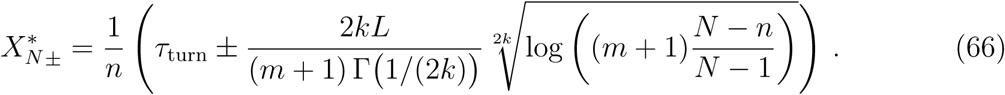

Similarly to *τ*_min_ and 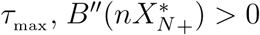 and 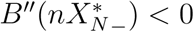, so from theorem 1, the unique ESS_N_ is

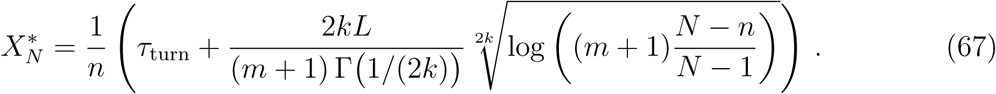

#### The curvature of the benefit function at the ESS_N_

Similar to equation (62), we have

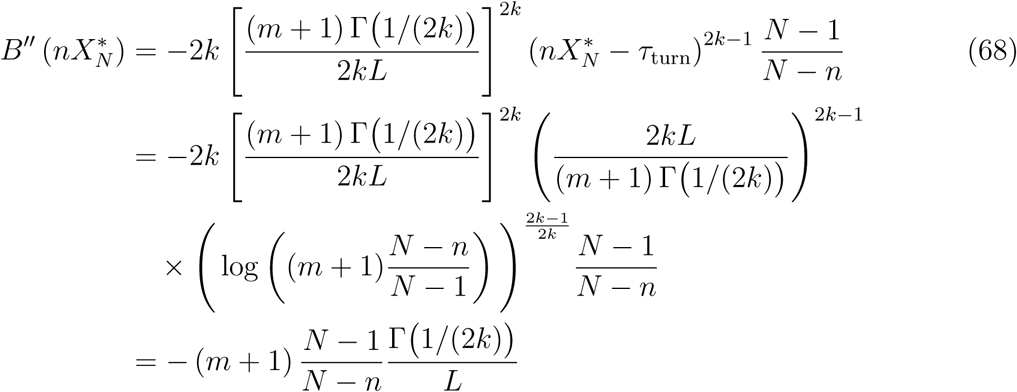

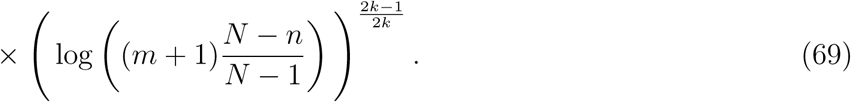

#### Condition for the fitness difference having a minimum when a single mutant defects and residents play the ESS

To guarantee that when a single mutant invades a population playing the ESS, the fitness difference has both a minimum and a maximum (as a function of the mutant strategy), we need the mutant contribution that minimizes fitness to be positive (or equivalently, the contribution of the nonfocal individuals—all residents—must be less than the minimizing total good *τ*_min_),

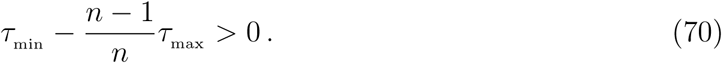

Using equations (59b) and (60), this is equivalent to *τ*_max_ *> n*Δ*τ*, or

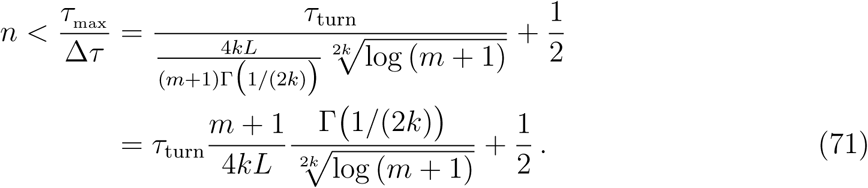

Rewriting this condition in terms of the horizontal asymptote *L*,

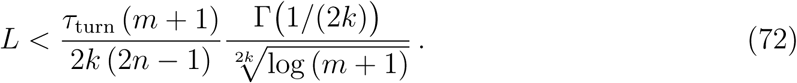

#### The payoff extrema difference

We now calculate the payoff extrema difference (PED), ΔΨ, that is, the difference between a mutant’s local minimum and maximum fitnesses when residents contribute the infinite-population ESS.

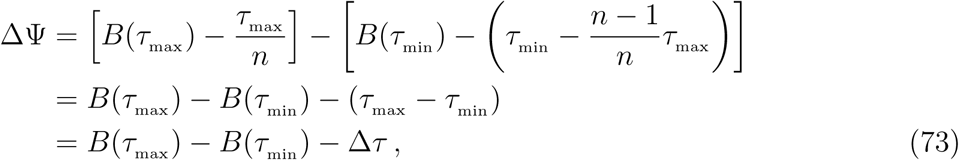

so using equations (60) and (63), we have

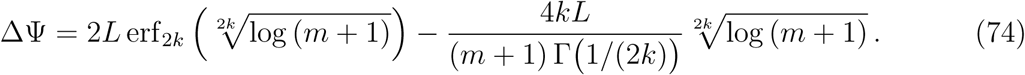

#### The mean fitness slope

To choose parameter values that generate a fitness difference with a distinct peak at the ESS (when residents play the ESS), we would like to find the mean fitness slope between the extrema, *i.e.*, the ratio of the PED, ΔΨ, and the distance between the fitness extrema as a function of our parameters. To that end, using equation (70), the distance between the fitness e trema is

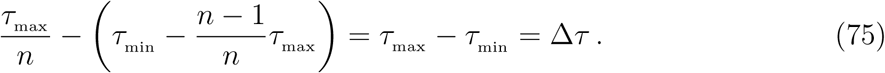

Equations (60) and (74) then yield

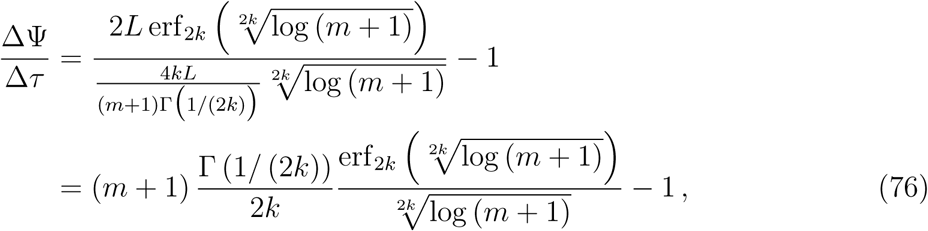

which only depends on the maximal marginal fitness, *m* (and the order of the generalized error function, 2*k*). Note also that using equation (48) and L’Hôpital’s rule (49),

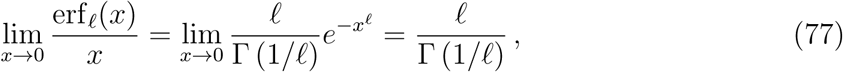

So

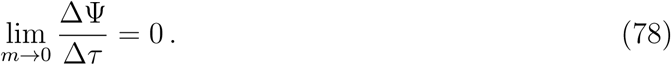

In addition, equation (49) implies that for any *x >* 0,

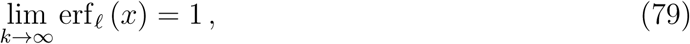

(because Γ(*x*) *→ ∞* as *x →* 0, and 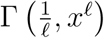 is bounded), and

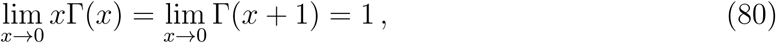

so we have

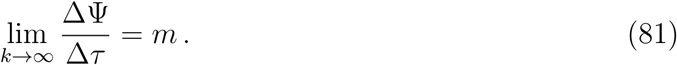

#### The ratio of ESSs in infinite and finite populations

Using equations (60), (61) and (67),

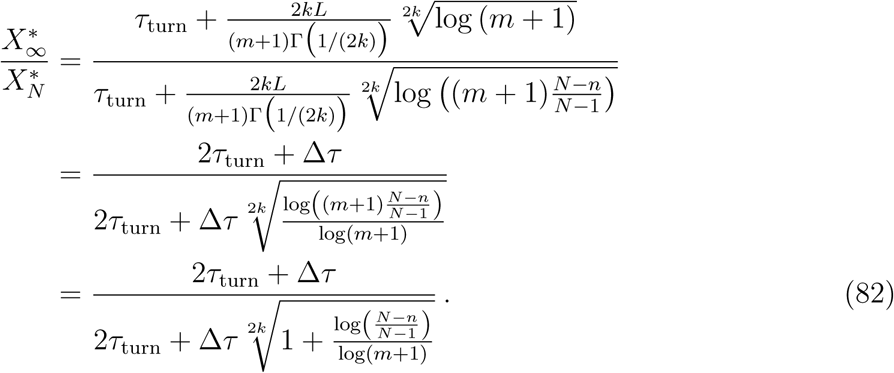

Rewriting the population size as *N* = *nG*,

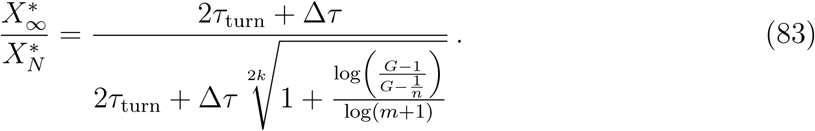

We see that the ratio 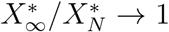 as *G → ∞* with *n* fixed. However, 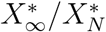 approaches a (finite) value greater than 1 as *n →∞* with *G* fixed (assuming 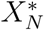 exist for all *N*; see inequality (15)).

### B.2 Sigmoid using standard error-function

In the special case *k* = 1 (*i.e.*, ℓ= 2), since 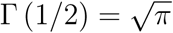, equation (47) reduces to

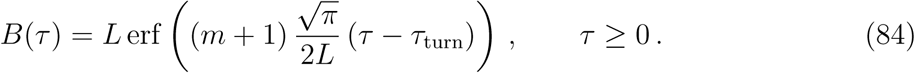

Then, setting *k* = 1 in equation (85) gives the maximizing and minimizing total goods,

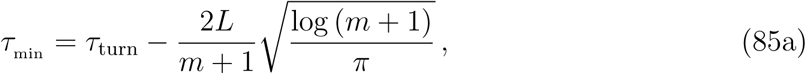

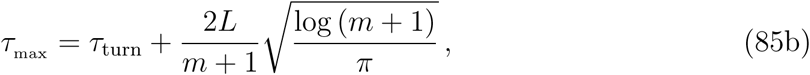

and the distance between the location of the fitness minimum and maximum is

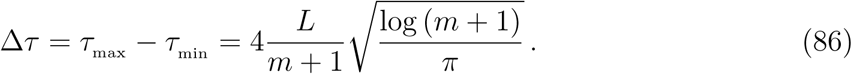

equation (61) then gives

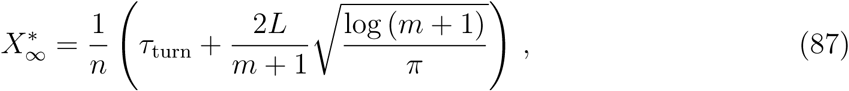

and equation (62) becomes

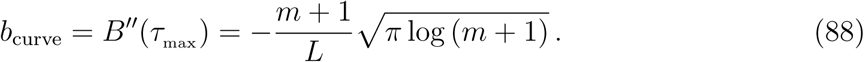

From equation (63),

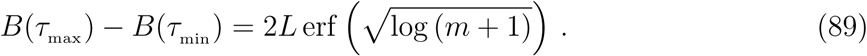

equation (67) gives the unique ESS_N_:

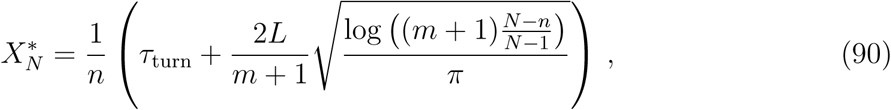

and equation (68) becomes

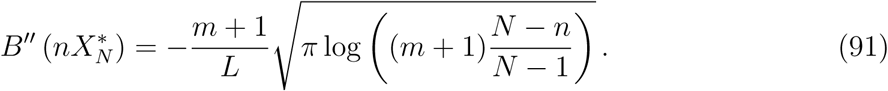

Condition (71), which guarantees that when a single mutant invade a population playing the ESS, the fitness difference has both a minimum and a maximum (as a function of the mutant strategy), reduces to

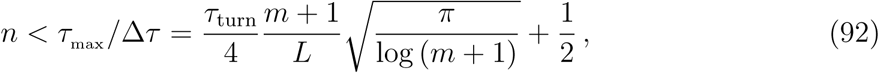

the PED, ΔΨ (equation (74)) becomes

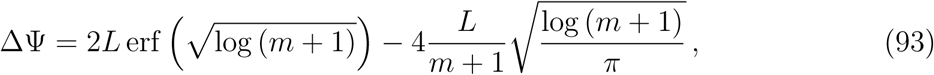

and the mean fitness slope (equation (76)) between the extrema reduces to

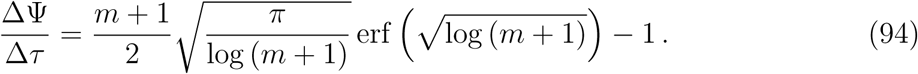

## Footnotes

∗ We assume throughout this paper that the strategy space is one-dimensional.

† In (11a), “generically” means excluding the unlikely possibility of singular strategies also being inflection points of *B*(*nx*); in (11b), it excludes the possibility of the marginal benefit *B*^*’*^ (*τ*) being constant in a neighbourhood of arg max *B*^*’*^ (*τ*).

‡ Condition (c) in the definition of the NSG (§5.1) implies that *m >* 0, so *N*_min_ is always well-defined in (13).

§ Note that *n*_c_ is always finite for a given population size, but when the number of groups *G* is fixed and larger than 1 + 1*/m*, then there is an ESS_N_ for any number of players *n*.

¶ More precisely, for a given maximal marginal fitness (*m*) and horizontal asymptote (*L*), *k* controls the distance between the benefit function’s inflection point (*τ*_turn_) and the total contribution at which the marginal benefit is half of its maximum.

\# equation (26) remains valid if individual fitnesses are obtained by averaging payoffs from an arbitrary (either fixed or random) number of rounds of the NSG, as long as groups are selected independently in each round.

‖ We need to avoid the situation in which singular strategy 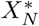 is also an inflection point of *B*(*nx*). This occurs when 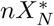 is both a critical point and an inflection point of *B* (*x*) *-* (*N -* 1) *x/* (*N - n*), which is generically *not* the case.

∗∗ For any positive integer *k*, Γ(*k*) = (*k -* 1)!.

